# On the limits of 16S rRNA gene-based metagenome prediction and functional profiling

**DOI:** 10.1101/2023.11.07.564315

**Authors:** Monica Steffi Matchado, Malte Rühlemann, Sandra Reimeiter, Tim Kacprowski, Fabian Frost, Dirk Haller, Jan Baumbach, Markus List

**Author notes:** Corresponding authors: Markus List.

## Abstract

Molecular profiling techniques such as metagenomics, metatranscriptomics or metabolomics offer important insights into the functional diversity of the microbiome. In contrast, 16S rRNA gene sequencing, a widespread and cost-effective technique to measure microbial diversity, only allows for indirect estimation of microbial function. To mitigate this, tools such as PICRUSt2, Tax4Fun2, PanFP and MetGEM infer functional profiles from 16S rRNA gene sequencing data using different algorithms. Prior studies have cast doubts on the quality of these predictions, motivating us to systematically evaluate these tools using matched 16S rRNA gene sequencing, metagenomic datasets, and simulated data. Our contribution is threefold: (i) using simulated data, we investigate if technical biases could explain the discordance between inferred and expected results; (ii) considering human cohorts for type 2 diabetes, colorectal cancer and obesity, we test if health-related differential abundance measures of functional categories are concordant between 16S rRNA gene-inferred and metagenome-derived profiles and; (iii) since 16S rRNA gene copy number is an important confounder in functional profiles inference, we investigate if a customised copy number normalisation with the rrnDB database could improve the results. Our results show that 16S rRNA gene-based functional inference tools generally do not have the necessary sensitivity to delineate health-related functional changes in the microbiome and should thus be used with care. Furthermore, we outline important differences in the individual tools tested and offer recommendations for tool selection.

## 1. Introduction

Microbial profiling through next-generation sequencing techniques has linked microbial composition to disease [1–3]. A common microbiome profiling approach is 16S rRNA gene sequencing which captures information on taxonomy and microbial diversity [4,5]. While accessible and cost-effective, 16S rRNA gene sequencing cannot directly provide insights into the biological processes in which microbes are engaged [6,7]. Taxonomic assignment of 16S rRNA gene sequences can give information about the identity of the microbes present in a community. Still, it does not necessarily provide information about their functional capabilities or the functional genes and pathways they possess. Microbial species can have different functional capabilities, even if they are taxonomically similar. For example, different strains or species of bacteria may have different metabolic pathways, allowing them to utilize different nutrients or tolerate changing environmental conditions [8,9]. Therefore, predicting functional profiles from 16S rRNA gene data relies on statistical inference and machine learning algorithms that use taxonomic information as a proxy for functional potential. These insights into microbial genes and alterations in pathways are crucial to understanding the role of the microbiome in health and disease [10].

In contrast to 16S rRNA gene sequencing, shotgun metagenome sequencing (MGS) provides rich information on functional genes, which can be translated to the pathway level using tools such as HUMAnN3 [11], Kraken [12], or MGS-FAST [13]. However, the considerably higher cost of MGS hinders its application in clinical research, particularly in large cohort studies, where adequate statistical power is essential for robustly detecting meaningful differences [14,15]. Additionally, MGS can be challenging for low biomass samples or samples that are dominated by non-microbial DNA [16,17].

To overcome the lack of functional information in 16S rRNA gene profiles, tools such as Phylogenetic Investigation of Communities by Reconstruction of Unobserved States (PICRUSt2) [18], Tax4Fun2 [19], Pangenome-based Functional Profiles (PanFP) [20], Piphilin [21], COWPI [22] and metagenome-scale models (MetGEMs) toolbox [23] attempt to predict abundances of functional genes based on recorded genomic information available in the Kyoto Encyclopedia of Genes and Genomes (KEGG) database [24] or based on genomic models [25– 27]. PICRUSt2 is a widely used prediction tool employing a hidden state prediction algorithm to infer functions from 16S rRNA gene phylotypes [18]. In contrast, Tax4Fun2 uses sequences within a defined similarity cutoff of reference sequences [19]. PanFP is based on a functionally annotated pan-genome reconstruction which is then weighted with the microbial abundance observed in a given sample [20]. Even though the accuracy of these tools has been validated, they are generally limited by the quality of the reference genomes and annotation, which suffer from ambiguous or missing coding regions [19,22]. To overcome this limitation, MetGEM introduced a generalised genome-scale model, where metagenome-scale networks are constructed using the AGORA collections [25] and the Human Microbiome Project (HMP) [28].

Another bias in amplicon sequencing is the number of 16S rRNA gene copies which varies considerably, confounding the abundance prediction [29,30]. Several tools have recently been developed for predicting genomic copy numbers using phylogenetic methods [18] and sequenced genomes [31]. The Ribosomal RNA Operon Copy Number Database (rrnDB) [32] offers accurate and well-annotated information on rRNA operon copy numbers among prokaryotes. Each entry (organism) in rrnDB contains detailed information linked directly to external websites, including the Ribosomal Database Project [33], GenBank [34], PubMed, and several culture collections.

For an accurate analysis and interpretation of functional profiles from 16S rRNA gene profiles, it is crucial to understand the limitations of these tools compared to more comprehensive profiling techniques such as MGS. Furthermore, it is essential to offer potential users guidelines on tool selection. To date, there are only a few reviews evaluating the performance of functional inference tools in different environments [35–38]. For example, Djemiel *et al*. [36] used a text-mining approach to review 100 published articles on functional profiles. They discussed limitations, including the lack of reference genomes, especially for soil ecosystems. Sun *et al*. [37] investigated the performance of three commonly used metagenome inference tools (PICRUSt, PICRUSt2, and Tax4Fun) across seven different datasets for which matched 16S rRNA gene and MGS data was available. Even though MGS data is not a truly gold standard of functional activity in the microbiome, it can serve as a gold standard for benchmarking prediction methods that infer functionality from 16S rRNA gene profiling to approximate MGS data. A striking finding of Sun *et al*. was that inferred abundances showed a high Spearman correlation between 16S-inferred and MGS-derived gene abundances even when sample labels were permuted. This highlights that functional profiles do not differ as much as changes in taxonomic composition suggest and that correlation is not a suitable performance measure to assess the performance of functional inference tools. The authors further showed that investigating a specific contrast, i.e. the difference between two groups with a Wilcoxon rank-sum test yielded p-values that were moderately correlated between 16S-inferred and MGS-derived genes and very lowly correlated after sample permutation. In their study, the authors concluded that functional inference tools worked well for humans rather than for other organisms and environmental samples and that core functions are better represented than niche-specific functions. However, the contrasts that were investigated were related to geography, where microbial composition and function are known to differ considerably [39].

It remains an open question if functional inference tools are also suited for more subtle contrasts related to human health. Sun *et al*. [37] could not detect any performance differences between the tested methods, suggesting that a more comprehensive benchmark is needed to recommend tool selection guidelines and establish the limits of metagenome-inferred tools in human disease research. Hence, we consider the most widely used metagenome inference tools PICRUSt2 [18], PanFP [20], Tax4Fun2 [19], and MetGEM [23] in a systematic benchmark. We omit Piphillin [21] as it is only available as a web server (the command line version is in the testing stage). We test the reliability of the inference tools using matched 16S rRNA gene sequencing and MGS human datasets both simulated as well as obtained from different cohorts, including Cooperative Health Research in the Augsburg Region cohort (KORA) (type 2 diabetes) [40], The Food Chain Plus (FoCus), PopGEN (obesity) [41], colorectal cancer (CRC) [42]. We use these data to examine the concordance of health-related functional changes in the microbiome between 16S-inferred and metagenome-derived profiles across commonly used software tools. We further explore the potential impact of 16S copy number as a confounding factor in functional profiles and investigate if a customised copy number normalisation using the rrnDB database could improve the results. In summary, our results support that current metagenome inference tools lack the sensitivity to identify health-related functional changes in the microbiome.

## 2. Materials and methods

### 2.2. Simulated dataset

We evaluated the performance of the functional inference tools using simulated metagenomic samples downloaded from the 2nd Critical Assessment of Metagenome Interpretation (CAMI) Challenge [43,44]. CAMI includes a set of simulated metagenomic datasets with known ground truth, allowing for objective comparison of tool performance. We retrieved 40 simulated metagenomic datasets from CAMI, which represent typical microbiomes from four human body sites such as gastrointestinal tract (GI) (n=10), skin (n=10), oral cavity (n=10), and airways (n=10).

### 2.2. Population-based cohorts

We also used real datasets from four population / patient cohorts. Naturally, these suffer from the usual technical bias and variability due to differences in sequencing platforms, protocols, and sample preparation methods. This allows us to assess the impact of those in comparison to well-controlled simulated data. We selected four cohorts with paired 16S rRNA gene sequencing and MGS data for functional profiling analysis. The CRC dataset was downloaded from the Sequence Read Archive (SRA) [45] under project number PRJEB6070. Datasets of FoCus, KORA, and PopGen are under controlled access due to the informed consent given by the cohort study participants and are available upon request from https://epi.helmholtz-muenchen.de/ and n https://portal.popgen.de, respectively. For the prospective KORA cohort (2018), sample preparation and sequencing of the V3V4 region in paired-end mode on an Illumina MiSeq was performed by the ZIEL – Core Facility Microbiome in Freising. For the FoCus cohort and PopGen cohort, a detailed overview is given in Table 1. The overall workflow is represented in Figure 1.

**Table 1:** Basic characteristics of population cohorts used in the benchmark study.

| Cohort | Geographic location | Sample size | Disease Focus | 16S rRNA gene region | Project URL | Ref. |
| --- | --- | --- | --- | --- | --- | --- |
| KORA | Augsburg, Bavaria, Germany | 60 | Diabetes vs Healthy | V3V4 | https://epi.helmholtz-muenchen.de/ | [40] |
| Focus | Kiel, Schleswig-Holstein, Germany | 101 | Healthy vs Obese | V3V4 | https://portal.popgen.de | [46] |
| Popgen | Kiel, Schleswig-Holstein, Germany | 86 | Healthy vs Obese | V3V4 | https://portal.popgen.de | [41] |
| CRC | Heidelberg, Germany | 182 | CRC vs Healthy | V4 | https://www.ncbi.nlm.nih.gov/bioproject/?term=PRJEB6070 | [42] |

**Figure 1:**
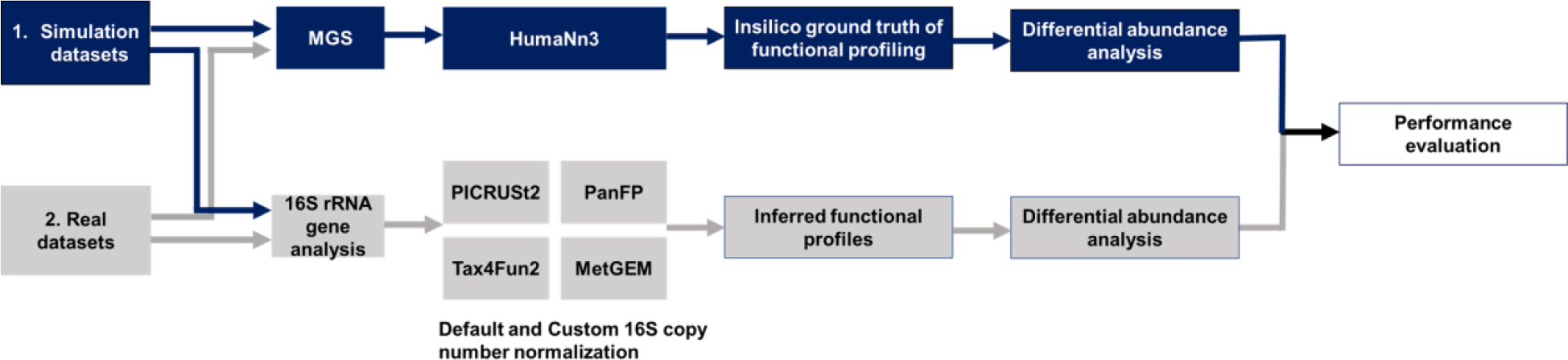
Overall benchmarking workflow to compare and evaluate the performance of functional inference tools from 16S rRNA gene sequences or metagenomics. First, paired WGS-16S rRNA gene datasets are selected for comparison. Here, simulated data (a) or real datasets (b) are considered. In the next step, we retrieve the functional profiles using Humann3 for the MGS datasets and four different functional inference tools (PICRUts2, Tax4Fun2, PanFP, and MetGEM) for 16S rRNA gene sequencing datasets. A Wilcoxon rank-sum test was performed to compare the KO terms resulting from the two methods.

### 2.3. Functional profiling of metagenomics data

For public datasets, sequence quality control and removal of human reads were performed using KneadData 0.7.4. [47] Bacterial gene abundances were calculated with HUMAnN3 (version 3.0) [11] using nucleotide-based alignment against the Chocophlan database [48]. Gene abundance tables were then grouped using the *uniref90_ko* command and relative abundance was calculated using the *humann_renorm_table* command in the HUMAnN3 pipeline. We followed the same strategy for the simulation datasets w.r.t. metagenome functional profiling, whereas functional profiling of 16S rRNA gene was done in two steps: First, 16S rRNA full length reads were filtered using SILVA [49] as a reference database. Once the 16S rRNA gene sequences were obtained, functional profiles using all four tools PICRUSt2, Tax4Fun2, PanFP, and MetGEM, were inferred as described below. We then compared the functional profiling from 16S rRNA gene analysis with the metagenome profiles and tested their agreement using functional diversity as mentioned in parallel-meta suite [50,51].

### 2.4. Functional inference of 16S rRNA gene sequencing data

Raw 16S rRNA gene reads were processed using DADA2 (1.26) [52] to obtain an amplicon sequence variants (ASV) abundance table. The representative sequences were then used as input for functional inference tools. We screened the literature for commonly used 16SrRNA gene-based functional inference tools and selected PICRUSt2 (v2.5.1), Tax4Fun2, PanFP and MetGEM for the benchmark analysis. PICRUSt2 and Tax4Fun2 are currently the most frequently used tools followed by PanFP. MetGEM represents a more recent development and has thus far not been benchmarked. The workflow for each tool is described in the corresponding methods subsection.

#### 2.4.1. PICRUSt2

ASV abundance tables with representative sequences were given as input for PICRUSt2 [18]. PICRUSt2 was executed with the default cutoff for the Nearest Sequenced Taxon Index (NSTI) of 2.0. The values in the copy-normalized ASV table produced by PICRUSt2 were multiplied by the expected 16S-rRNA copies per taxon (also output by PICRUSt2) to obtain an untransformed ASV table of contributing sequences. The totals of each column were then recorded as the total number of sequences used by PICRUSt2 in each sample.

#### 2.4.2. Tax4Fun2

Tax4Fun2 was run with the default pre-calculated files and a parameter of 99% similarity in its *BLASTn* command. Tax4Fun2 outputs relative abundances by default, so no transformation was required after producing the prediction profile. An adjustment was made to the Tax4Fun2 script *make_FunctionalPredictions()* to output the ratio of used sequences instead of the default ratio of unused sequences. Note that at the time of manuscript submission, Tax4Fun2 was no longer available at https://github.com/bwemheu/Tax4Fun2.

#### 2.4.3. MetGEM

The MetGEM toolbox is based on the Genome-scale models for inferring the metagenomic content from 16S rRNA gene sequences with an overall focus on annotating the metabolic function of the human gut microbiome. ASV abundance tables along with corresponding taxonomic groups were given as input. Default models such as k_core and e_core were chosen to predict KEGG orthologs (KO) and Enzyme Nomenclature (EC) abundances, respectively, which were previously shown to provide good estimation in typical situations [23]. Since MetGEM does not provide pathway abundances, the output from MetGEM was subjected to the PICRUSt2 pathway prediction step.

#### 2.4.4. PanFP

PanFP [20] is based on the pangenome reconstruction of a 16S rRNA gene operational taxonomic unit (OTU) from known genes and genomes pooled from the OTU’s taxonomic lineage. PanFP includes complete genomes of prokaryotes with their full taxonomic lineages obtained from the National Center for Biotechnology Information (NCBI) [53] and mapped KO terms to proteins using the cross-reference ID mapping between NCBI RefSeq [54] and UniProt provided by UniProt KnowledgeBase (UniProtKB) [55]. For each OTU, PanFP builds a pangenome by making a superset of all genes present in organisms pooled from the dataset of prokaryote genomes in the given OTU’s taxonomic lineage. Therefore, OTUs with the same taxonomic lineages have the same pangenome. Then, PanFP derives a functional profile of the pangenome by accumulating functional compositions in the superset. An OTU-sample table is converted into a lineage-sample table by summing the frequencies of OTUs with the same lineage in a sample. Finally, the function-sample table is derived by combining functional profiles of lineages with weights corresponding to the lineage abundance in the sample.

### 2.5. Customized normalization using rrnDB

To test the effect of copy number normalization on the functional profiling, we repeated the entire workflow described above, replacing the gene copy number normalization step by obtaining the copy numbers from rrnDB. We processed rrnDB with the PICRUSt2 *place_seq*.*py* command (see https://github.com/mruehlemann/16s_cnv_correction_databases for details) and used the resulting abundance tables as input for other tools by skipping their built-in copy number normalization steps.

### 2.6. Validating inference tools with shotgun metagenomic sequencing data

We used two methods to evaluate the consistency and accuracy of functional inference tools. It should be noted that a direct comparison of functional profiles inferred with all four tools is virtually impossible due to several changes in the KEGG Orthology since PICRUSt2, Tax4Fun2, PanFP, and MetGEM were developed. Hence, inferred functional profiles as well as those obtained by metagenomic shotgun sequencing were converted to relative abundances prior to comparison. For metagenomics, the counts were converted into relative abundances using the *renorm table* function in HUMAnN 3.0. We analyzed the Spearman correlation between inferred gene composition and those from metagenome sequencing. Only functions present in the metagenomic profile and in the inferred profile were considered in each comparison.

### 2.7. Differential abundance analysis between cases

As proposed by Sun *et al*. [37], differential analysis is better suited than correlation analysis to assess whether metagenome inference tools are able to detect biological variation between samples. We thus evaluated the performance of inference tools in predicting biological functions using the differential abundance testing method. To do so, we removed KO terms with low prevalence (< 5% of samples) and retained overlapping KO and pathway terms between the prediction tool and MGS for differential analysis. We performed a Wilcoxon rank-sum test of the two groups (disease versus control) in each dataset for both default and customized normalization.

Using simulated datasets, we compared the KO abundance between GI versus skin and also GI versus airways to test the accuracy of the tools. Similarly, using the CRC dataset, we compared the KO abundance between cancer (n=41) and healthy tissues (n=50). Using the Popgen and FoCus datasets, we compared the differentially abundant KO terms between obese and healthy samples. Patients with a body mass index (BMI) > 30 kg/m^2^ were considered obese. For the KORA dataset, differentially abundant KO terms were tested between type 2 diabetes and healthy controls. The Wilcoxon rank-sum test was applied to each cohort to test the difference between groups for MGS and prediction results and inferred KO abundance and significant KO terms with a p-value < 0.05 were extracted. Overlapping significant KO terms between MGS and inference tools were extracted and compared to evaluate the performance of inference tools using the F1 score, recall, and precision metrics.

## 3. Results

### 3.1. Results for simulated data show that 16S-based functional inference can uncover functional differences between diverse environments

Simulated datasets provide a controlled environment for testing the performance of tools and can be constructed free of bias and variability due to differences in sequencing platforms, protocols, and sample preparation methods. Hence, we used this setup to show how close functional inference tools can be expected to approximate metagenome-derived functional profiles under optimal conditions. PCA was used to compare the performance of four different tools in predicting the functional potential of a microbial community based on 16S rRNA gene sequencing data (Figure 2). Based on the results of the PCA analysis, it was observed that PICRUSt2 and PanFP, using either default or customized normalization, were embedded more closely to the metagenomic profiles in the PCA space. This suggests that these tools were more accurate in predicting the functional potential of the microbial community, based on the 16S rRNA gene sequencing data. In contrast, Tax4Fun2 and MetGEM showed a high discrepancy from their corresponding metagenomes. This indicates that these tools may not be as accurate in predicting the functional potential of the microbial community based on 16S rRNA gene data.

**Figure 2:**
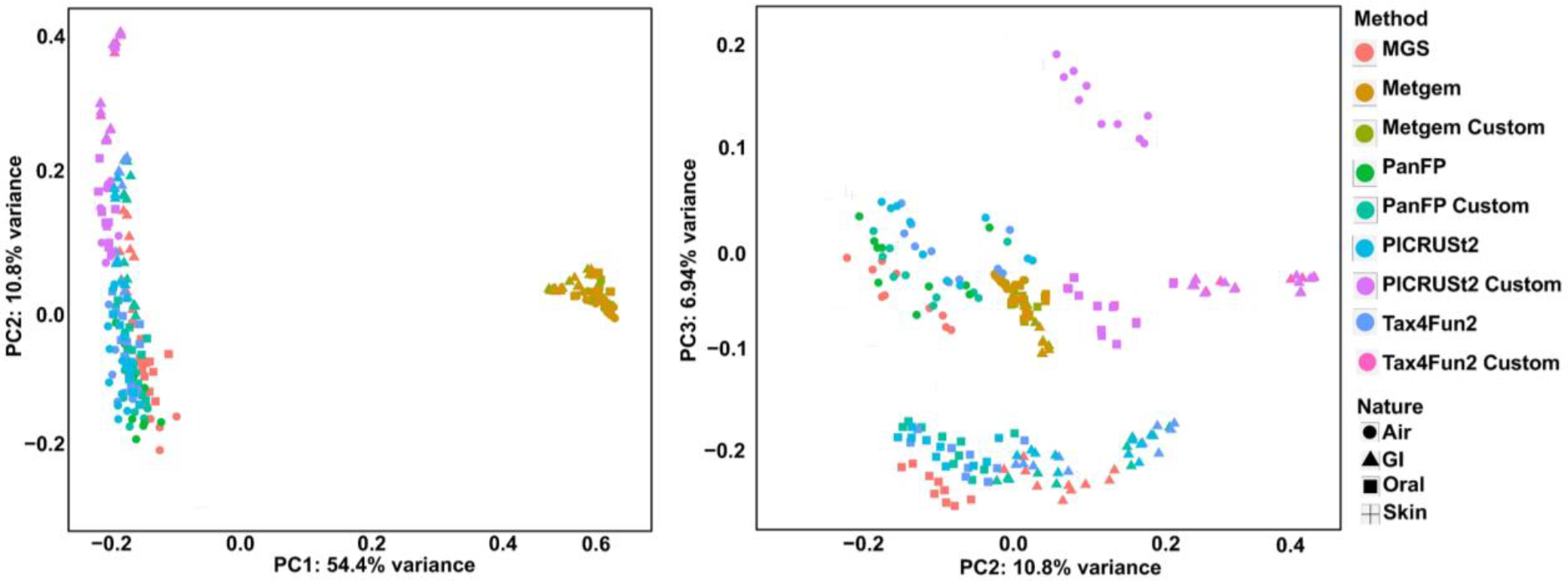
Principal Coordinates Analysis (PCoA) for functional profiling of simulated datasets for both functional inference tools and metagenome analysis. Using simulated datasets from different environments, inference tools such as PICRUSt2 and PanFP produce functional profiles overall similar to the ground truth metagenome functional profiling of HUMAnN3, whereas functional profiles from Tax4Fun2 and MetGEM are clustered far from the ground truth.

We further conducted differential abundance tests using KO terms detected in simulated MGS and 16S rRNA gene sequencing data. We focused on the comparisons of GI and oral cavity as well as GI and airways where we expected to find considerable variation in the functional profiles due to their distinct microbial communities and environmental conditions.

We considered significantly differentially abundant KO terms detected in MGS as ground truth and compared this list against significantly differentially abundant KO terms reported by functional inference tools, allowing us to calculate F1 score, recall and precision. In the GI vs. oral cavity comparison, PICRUSt2 and Tax4Fun2 displayed a moderately higher F1 score (ranging from 0.5 to 0.8) and recall compared to low-performing tools PanFP and MetGEM. In the GI vs airways comparison, PICRUSt2 again performed well with higher recall and F1 scores followed by MetGEM, Tax4Fun2, and PanFP. Notably, all tools displayed low precision (Figure 3). This suggests that even in the simulated datasets, functional inference tools tend to produce a large number of false positive results.

**Figure 3:**
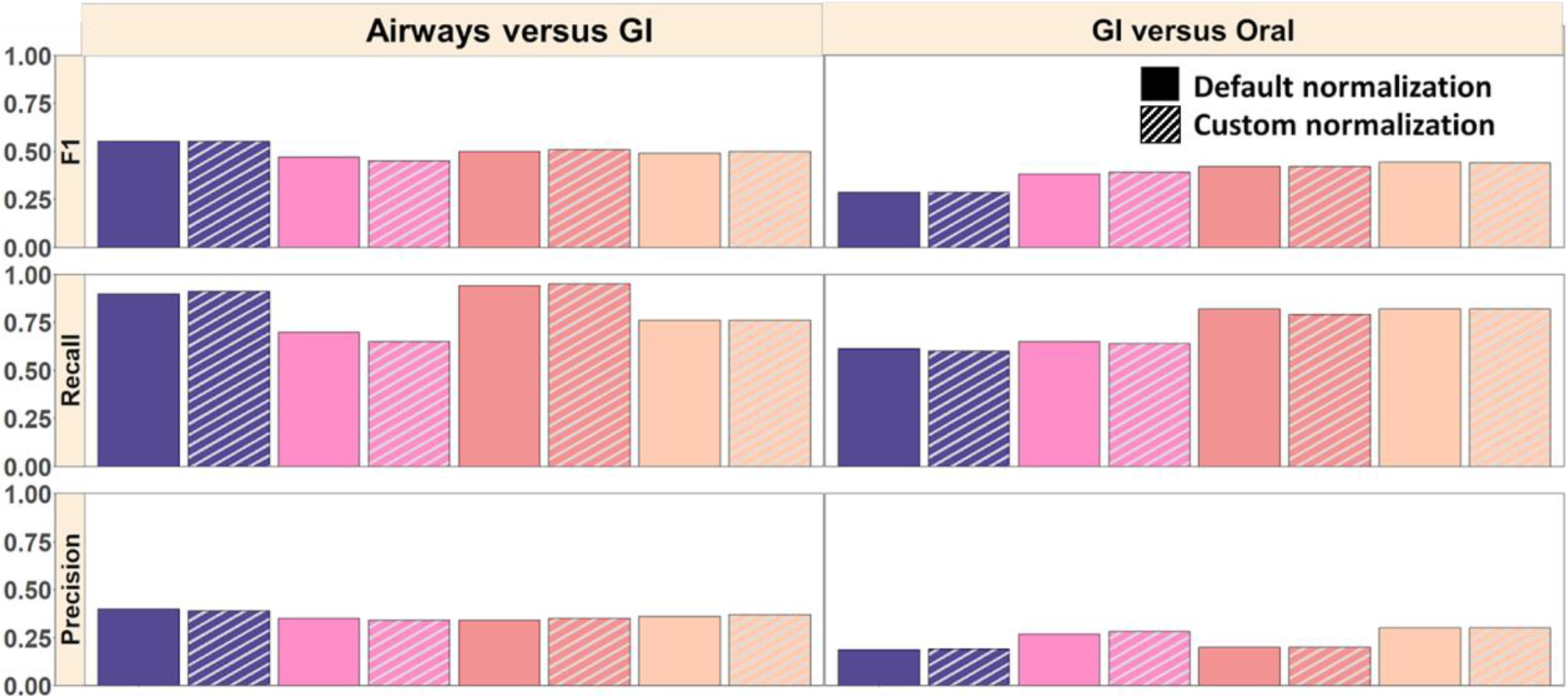
Comparison of significantly differentially abundant KOs between inferred and MGS results. F1 score, recall, and precision scores are reported for each category compared to the MGS data also between default and customized normalization.

### 3.2. Functional inference tools show good performance in predicting the presence of KO terms in real data

While simulated microbial datasets generated from 16S rRNA gene sequencing can be useful for benchmarking, they do not fully capture the complexity and variability of real-world microbial communities. One major limitation is the lack of amplification bias that occurs during PCR amplification [38,56,57] of the 16S rRNA gene. This bias can result in uneven amplification of different bacterial taxa, leading to overrepresentation or underrepresentation of certain microbial species in the resulting amplicon library and functional profile variation among phylogenetically related genomes. As a consequence, microbiome functional profiles inferred from the 16S rRNA gene can deviate from MGS-derived predictions. To evaluate the performance of analysis tools more accurately, we used paired MGS and 16S rRNA gene datasets from four different population cohorts (CRC, FoCus, Popgen, and KORA).

We confirmed the findings of an earlier study by Sun *et al*. [37], who reported that Spearman correlation values are not affected by label permutation and are thus not suited to robustly assess the performance of functional inference tools (Suppl. Fig S1). As an alternative, we consider here which KO terms are reported and compare them to those identified in MGS as ground truth. KO terms that are uniquely identified by 16S-based inference tools represent false positives, whereas missing KO terms in 16S functional inference tools that are reported by MGS represent false negatives, allowing us to compute F1 score, recall, and precision. Overall, Tax4Fun2, PICRUSt2, and PanFP showed similar performance in terms of F1 score and recall in contrast to MetGEM, which had already shown poor performance in the simulation study which is confirmed here (Figure 4).

**Figure 4:**
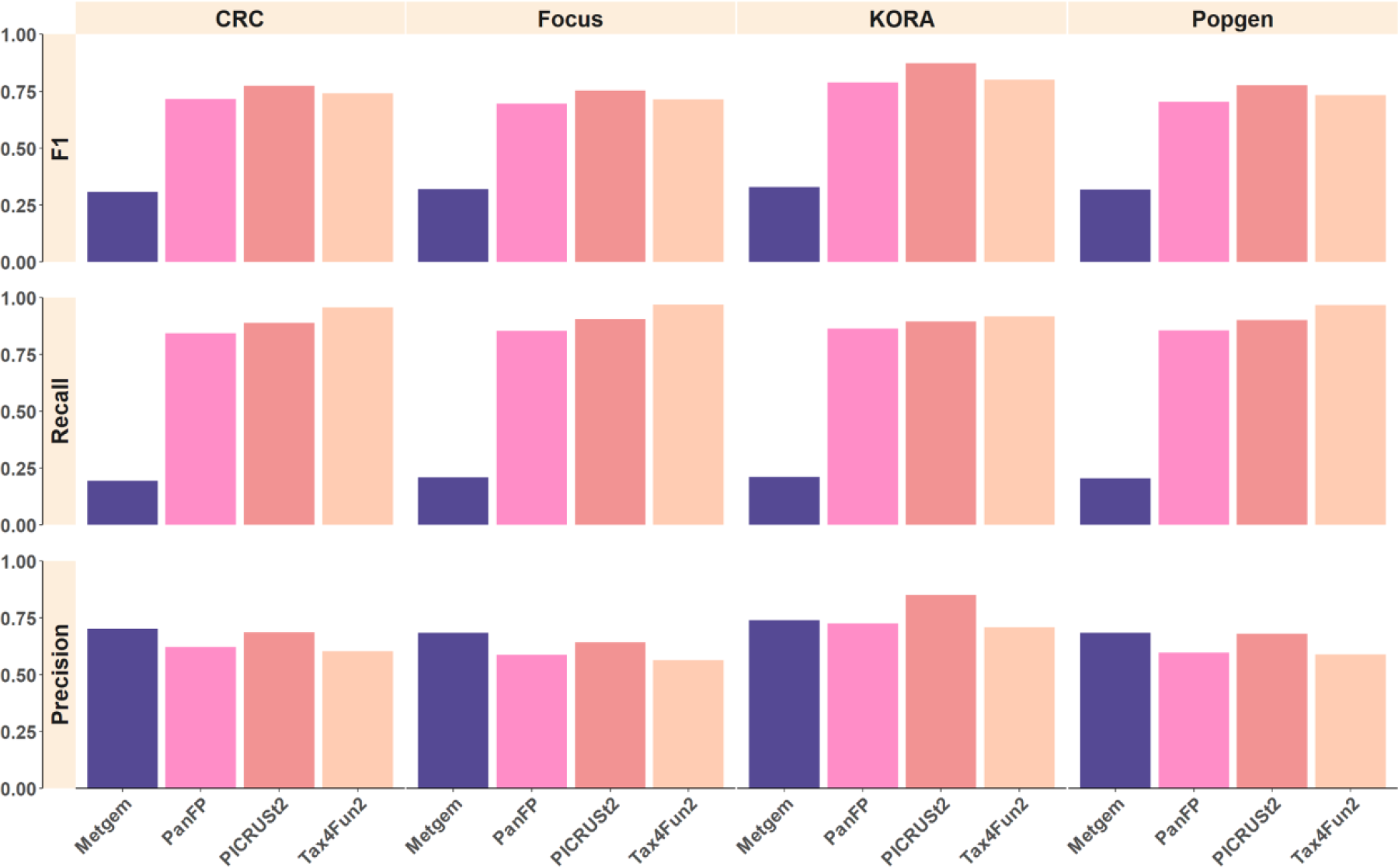
Comparison of detected KEGG terms between inferred metagenomes and MGS. F1 score, recall and precision are reported for each category compared to the MGS data. PanFP, PICRUSt2, and Tax4Fun2 show comparable and relatively consistent performance across datasets while MetGEM shows poor recall.

### 3.3. Functional inference tools show poor performance in predicting differentially abundant KO terms in real data

A more interesting question for biomedical research is, though, if these methods are accurate enough to pick up on health-related differences which may be subtle on a functional level. Hence, we performed a Wilcoxon rank sum test to identify differential pathways between healthy and disease groups, where we found that PICRUSt2, Tax4Fun2, and PanFP had varying degrees of overlap with MGS results. In the CRC cohort (Figure 5), PICRUSt2 shows the largest overlap of significant KO terms (n=654) with MGS results followed by Tax4Fun2 and PanFP. The number of overlaps of significant KO terms dropped significantly in other cohorts including KORA, FoCus, and Popgen (Suppl Fig 2-4). In the KORA cohort, Tax4Fun2 has the largest overlap of significant KO terms (n=112) followed by PICRUSt2 with custom normalization (n=108) and PICRUSt2 (n=66). PanFP with custom normalization (n=94) showed higher overlap than the PanFP with default normalization (n=13). In the Popgen and FoCus cohorts, the size of the overlaps dropped considerably, as only a few overlapping KO terms were found. PICRUSt2 with custom normalization and Tax4Fun2 showed a comparatively good overlap of significant KO terms in the Popgen and FoCus cohorts. PanFP and MetGEM showed poor overlap of KO terms across all cohorts.

**Figure 5:**
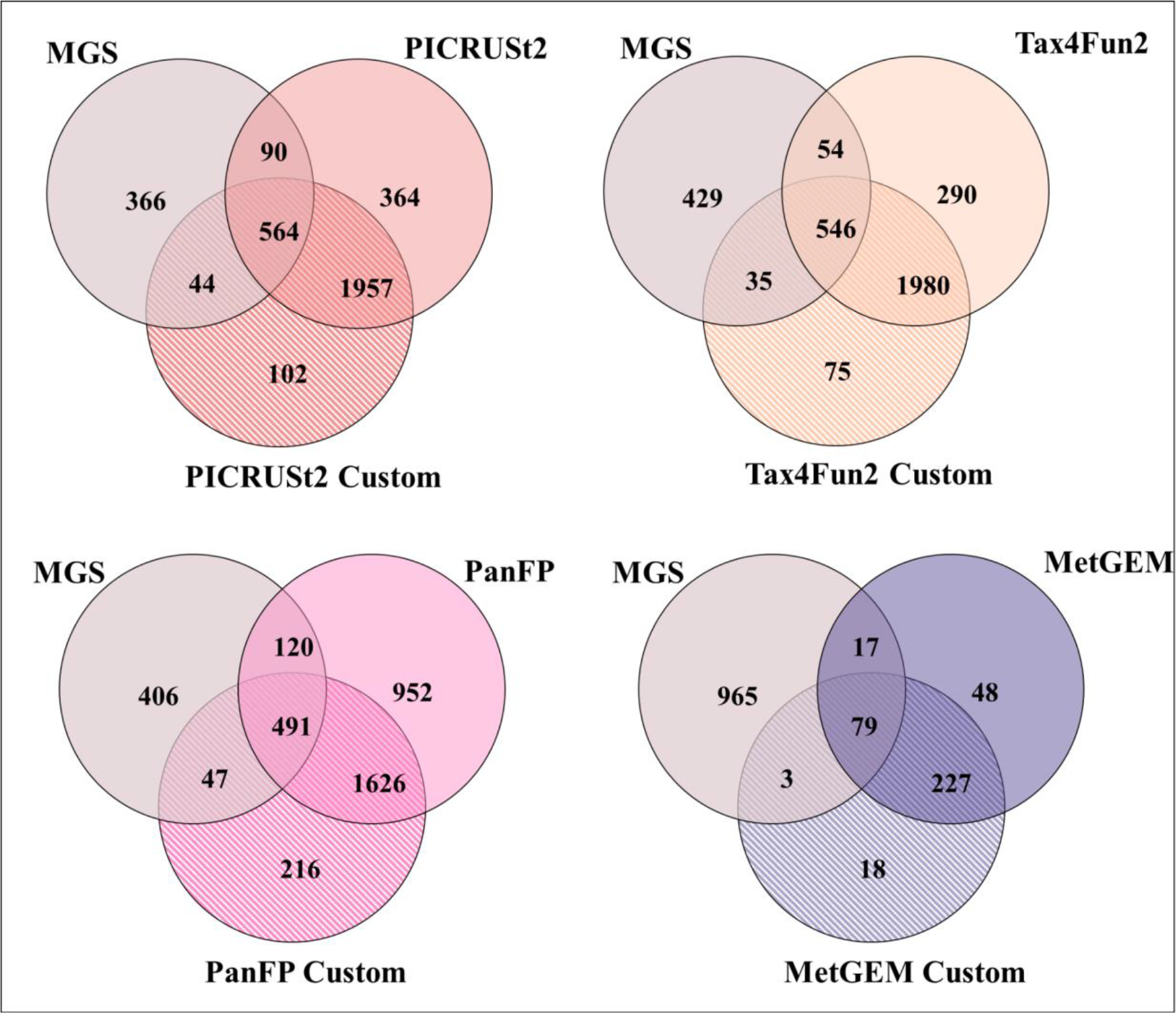
The overlap of significant KO terms between different functional inference tools and MGS results in the CRC cohort. The number of overlapping significant KO terms can be used as a quantitative indicator of the accuracy of the predictions.

Next, we calculated the F1 score, recall, and precision considering MGS as the ground truth for differentially abundant KO terms (Figure 6). Comparing the healthy and disease groups we obtained overall low F1 scores for all functional inference tools with markedly better performance in the CRC cohort due to a higher recall. The F1 scores of PICRUSt2, Tax4Fun, and PanFP were similar and could not be improved by customized copy number normalization, while MetGEM showed a slightly improved F1 score after copy number correction.

**Figure 6:**
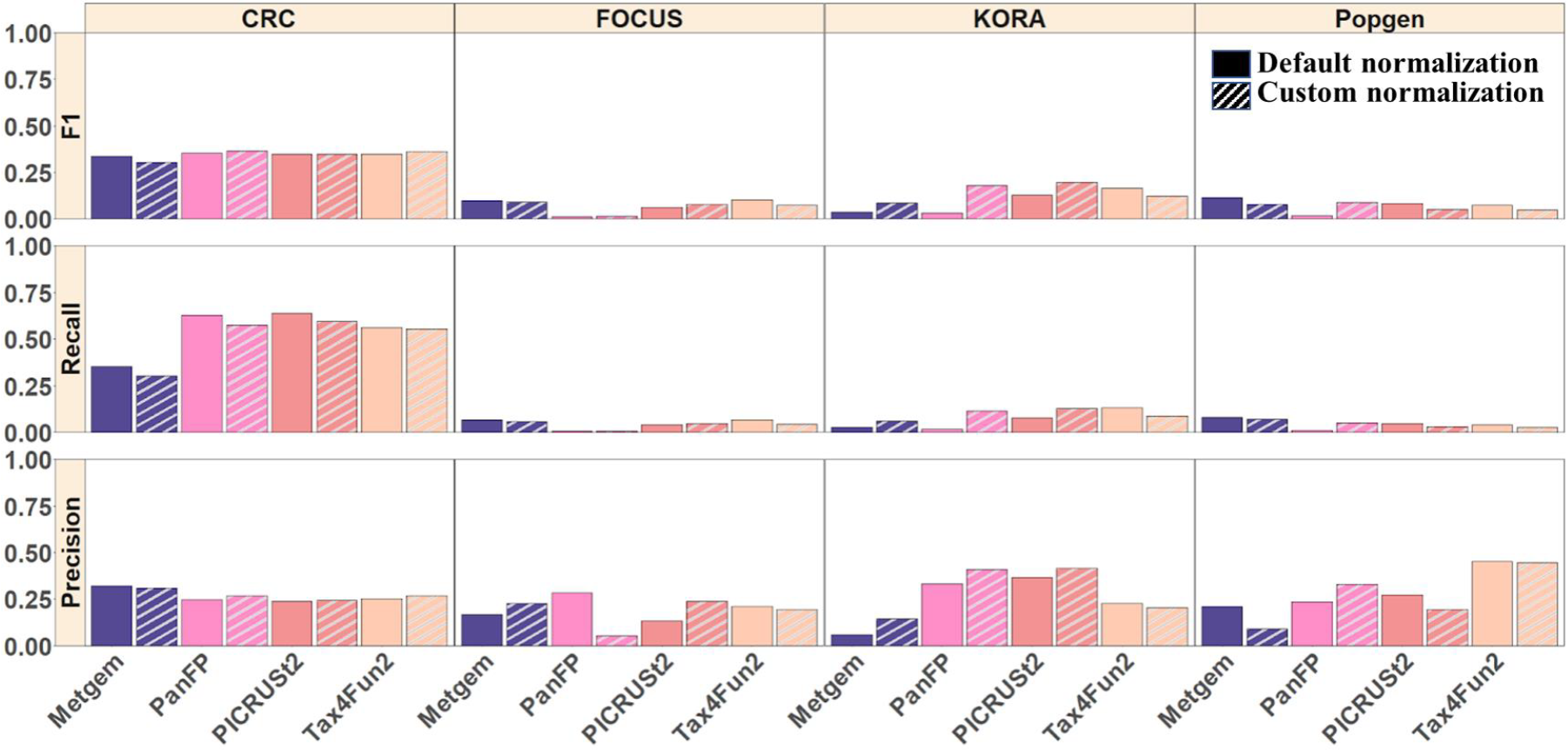
Comparison of significantly differentially abundant KO terms between inferred metagenomes and MGS. F1 score, recall, and precision are reported for each category compared to the MGS data.

We performed label randomization to assess the significance and robustness of the F1 score, recall, and precision values obtained from the original data (Figure 7). Randomization was performed only on the inferred functional profiles (not on the MGS data) at the sample levels and differential abundance testing was performed using the Wilcoxon rank sum test. Subsequently, the inferred p-values were compared with the p-values obtained from differential abundance testing performed on MGS. After label randomization, the groups we compare are random; hence, the differentially abundant pathways are not meaningful, and we obtain a baseline for our comparisons. We observed an improvement in recall in the CRC and a slight improvement in precision in the KORA and Popgen cohorts, whereas we observed equal or even worse performance in the FOCUS cohort.

**Figure 7:**
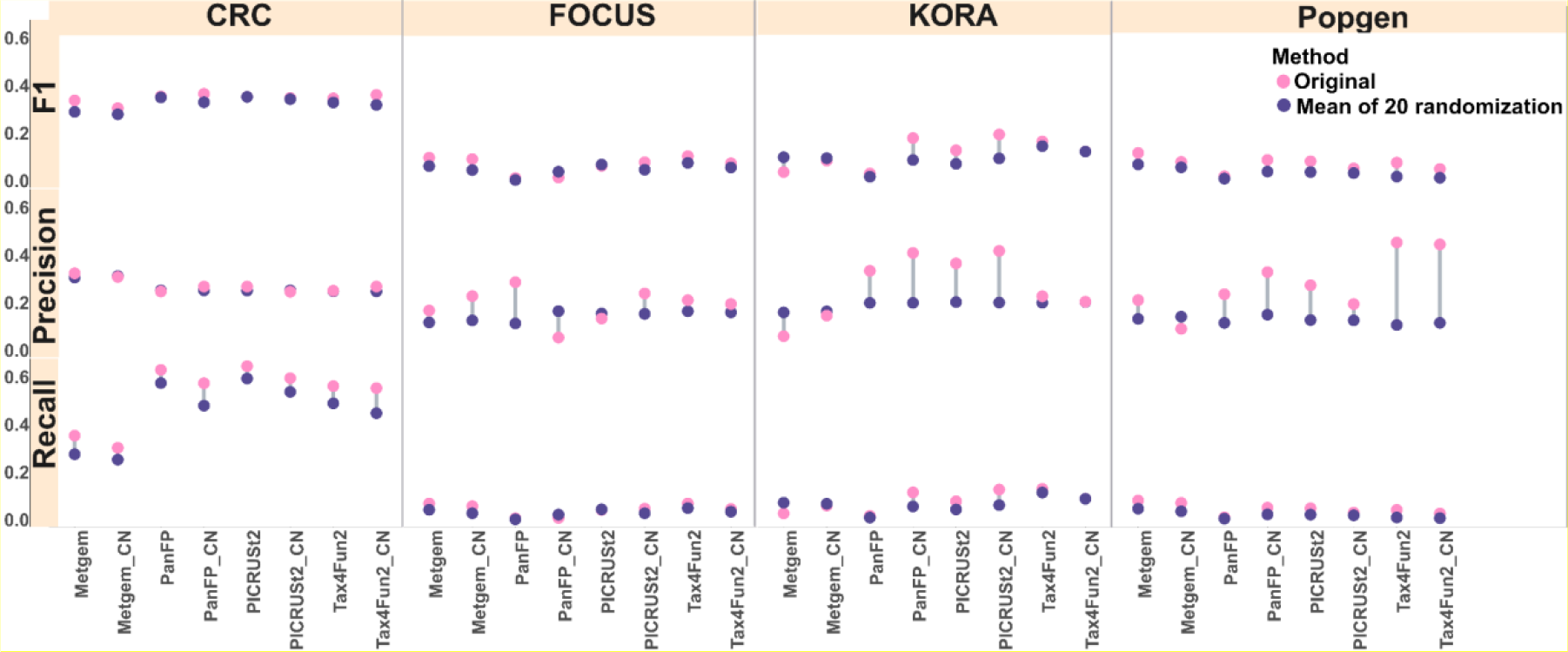
Comparison of significantly differentially abundant KO terms between inferred metagenomes and MGS was conducted before (original) and after 20 rounds of randomization. The randomization process was repeated 20 times to assess the significance and robustness of the F1 score, recall, and precision. A dumbbell plot was generated, depicting the mean value of the 20 randomizations against the original (ground truth) score.

### 3.4. In a targeted analysis many disease-relevant KO terms show differential abundance

We next focused on known disease-specific KO terms to assess if functional inference tools can detect differences in them. For the CRC cohort, we focused on KO terms classified under pathways in glycans biosynthesis and metabolism, lipid metabolism, carbohydrate and amino acid metabolism. These KO terms are reported to be enriched in CRC patients [58–62]. For the KORA cohort, we focused on KO terms classified under pathways in carbohydrate [63] and amino acid metabolism [64,65]. In diabetes, particularly in type 2 diabetes, impaired carbohydrate metabolism is a hallmark feature. The body’s ability to regulate blood glucose levels becomes compromised, leading to hyperglycemia. Insulin resistance and impaired insulin secretion contribute to this dysregulation [66]. Also, amino acids play essential roles in cellular metabolism and have implications for glucose homeostasis and insulin action. Perturbations in amino acid metabolism have been observed in diabetes [67,68]. Since the FoCus and Popgen cohort studies include both healthy and obese groups, we focused on KEGG functional categories such as carbohydrate metabolism, amino acid metabolic pathways, and sugar transport which have been reported as enriched in obesity [69,70]. We selected significant KO terms (p-value < 0.05) in the MGS data under the above-mentioned categories and compared them with KO terms obtained from the functional inference tools (Figure 8).

**Figure 8:**
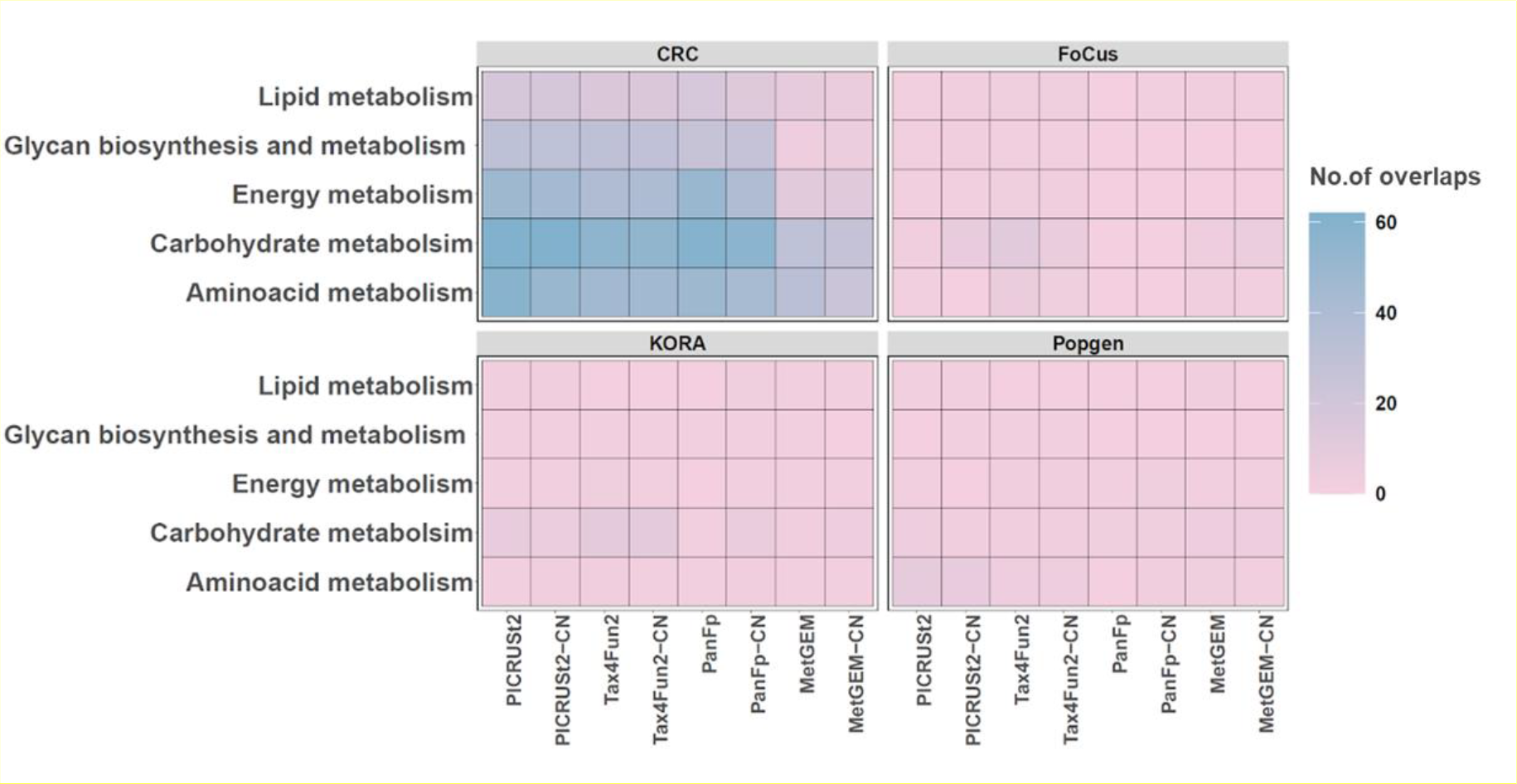
Heatmap representing the number of significant KO terms under each KEGG metabolic function between functional inference tools and MGS. Each cell in the heatmap corresponds to a specific combination of a disease-specific KO term and a tool. The colour of the cell indicates the number of overlapping KO terms detected by both MGS and the functional inference tools. Darker colour represents a higher number of overlapping KO terms, while paler colour indicates a lower number of overlaps.

Overall, PICRUSt2, Tax4Fun2 and PanFP showed better performance in the CRC cohorts. For the biosynthesis and metabolism of the glycans category, PanFP showed the largest overlap of significant terms with MGS followed by PICRUSt2 and Tax4Fun2 (default and customized normalization). There were only three overlapping significant KO terms between MGS and MetGEM. Similarly, the study compared KO terms in lipid metabolism and found that PICRUSt2 (default and custom normalization) showed a high number of overlaps with MGS, followed by Tax4Fun2 and PanFP (default and customized normalization).

In other cohorts, the performance of these tools, including PICRUST2 and Tax4Fun2, was poor. For example, in the KORA cohort and the pathway of carbohydrate metabolism, only PICRUSt2 (with default and custom normalization) showed a few overlapping significant KO terms with MGS. This indicates that the performance of other inference tools may be limited in predicting disease-specific KO terms related to carbohydrate metabolism in the context of the KORA cohort. Similarly, FoCus and Popgen cohorts showed very poor overlaps with MGS (individual abundance differences of KO terms for these functional categories were shown in Suppl Fig 7-9). In summary, these results show that functional inference tools have limited success in identifying disease-related functional activities.

## Discussion

Metagenome analysis is often the first choice for inferring functional relationships within the microbiome and between microbiomes and their ecosystem [71,72]. However, due to the lower costs, 16S rRNA gene profiling is still popular for studying microbial abundances and taxonomic profiles [73–75]. PICRUSt2, Tax4Fun2, PanFP, and MetGEM were developed to predict microbial functions from 16S rRNA gene sequencing datasets. They achieve this by utilizing reference genome databases such as KEGG and ancestral state reconstruction methods [18–20]. Each tool follows a different algorithmic approach and relies on different references, leading to a considerable prediction discrepancy. In this study, we sought to understand the limitations of these tools in human disease research.

We initially considered a simulated data set in which we generated full-length 16S profiles from simulated MGS data without the influence of technical and platform-related differences, e.g., related to PCR bias. In a global PCA analysis, PICRUSt2 and PanFP results closely matched the MGS-derived pathway abundances. Tax4Fun2 showed a shift from the ground truth but maintained the overall differences of the sample groups, whereas MetGem showed poor performance.

All metagenome inference tools except MetGEM showed overall good performance in predicting the presence of KO terms in real data. PICRUSt2 performed slightly better than other tools, possibly because the hidden state prediction increases the sensitivity to detect ubiquitous functions [38]. Another explanation could be that PICRUSt2 uses the most recent reference, i.e., the Integrated Microbial Genomes and Microbiomes (IMG/M) database, for predicting the functional content of microbial communities. On the other hand, MetGEM showed the lowest precision. One reason might be that the AGORA collections used as reference contain 818 genome-scale models (GEMs) from the human gut microbiome, which covered only 1,470 KO terms, 983 EC numbers across 226 genera, and 690 species in total [23,25].

Prior studies established a high Spearman correlation between inferred functional profiles and MGS-derived profiles that occurs even after label randomization [37]. In this study, we focused on the differential abundance of functional terms as a more challenging scenario. We observed poor performance across all functional inference tools, with PICRUSt2, PanFP and Tax4Fun2 showing comparable performance, whereas MetGEM showed very poor performance. All tools showed better performance in the CRC cohort compared to the other cohorts. We hypothesize that cancer-associated dysbiosis functional changes may be more pronounced than those in diabetes and obesity. Rather than looking at the differential abundances across the entire functional profile, one can also focus specifically on KO terms previously associated with the disease. Here, we observed significant differences in many of the KO terms also identified in MGS. This suggests that in contrast to a global analysis which is limited by false positives, a more targeted 16S functional profiling can pick up health-related differences to a certain extent.

A clear finding of this study is that functional profiles from 16S rRNA gene data are generally not suited to systematically pick up differences related to changes in human health in typical settings such as CRC, obesity or type-2 diabetes. Limitations in the performance of functional inference tools can be explained by an incomplete reference genome [76–78]. While functional inference tools pick up major differences, e.g., between ecological niches, they should not be used as a replacement for MGS in the study of human health. If researchers intend to produce functional profiles from 16S rRNA gene data for hypothesis generation they should be aware of these limitations and implement control strategies such as sample label randomization. Among the available tools, we recommend using PiCRUST2 and Tax4Fun, which appeared to be the most robust, followed by PanFP, which could be improved by introducing a customized copy number database.

## 5. Data availability

The repository at https://github.com/biomedbigdata/16S-rRNA-gene-Functional_benchmark_profiling includes the processed data files that can be used to re-generate the figures and findings in this paper. The accession codes for all sequencing data used in this study are listed in the Supplementary Methods.

## 6. Acknowledgements

Funded by the Deutsche Forschungsgemeinschaft (DFG, German Research Foundation) – Projektnummer 395357507 – SFB 1371 and 422216132. JB was partially funded by his VILLUM Young Investigator Grant nr.13154. JB received funding for this work through the FeMAI project, which is funded by the German Federal Ministry of Education and Research (BMBF) under grant number 01IS21079.

