## Supplementary material for "On the limits of 16S rRNA gene-based metagenome prediction and functional profiling": Supplment

### S1. Correlation is not a suitable performance measure for metagenome prediction tools

Sun *et al.* [37] previously investigated the performance of metagenome prediction tools and observed that the comparably high Spearman correlation values are not affected by label permutation. We could confirm these findings on five independent disease cohorts where PICRUSt2, Tax4Fun2 and PanFP achieved Spearman correlation values ranging from 0.65 to 0.75 **(Figure S1)** which did not significantly drop drastically after sample label permutation. MetGEM performed slightly worse than its competitors. Using rrnDB copy number normalisation, PICRUSt2, Tax4Fun2 and MetGEM did not show much improvement, while the performance of PanFP was raised to the level of the top-performing tool PICRUSt2. Since correlation analysis is not suited to robustly assess the performance of existing methods, alternative measures are needed.

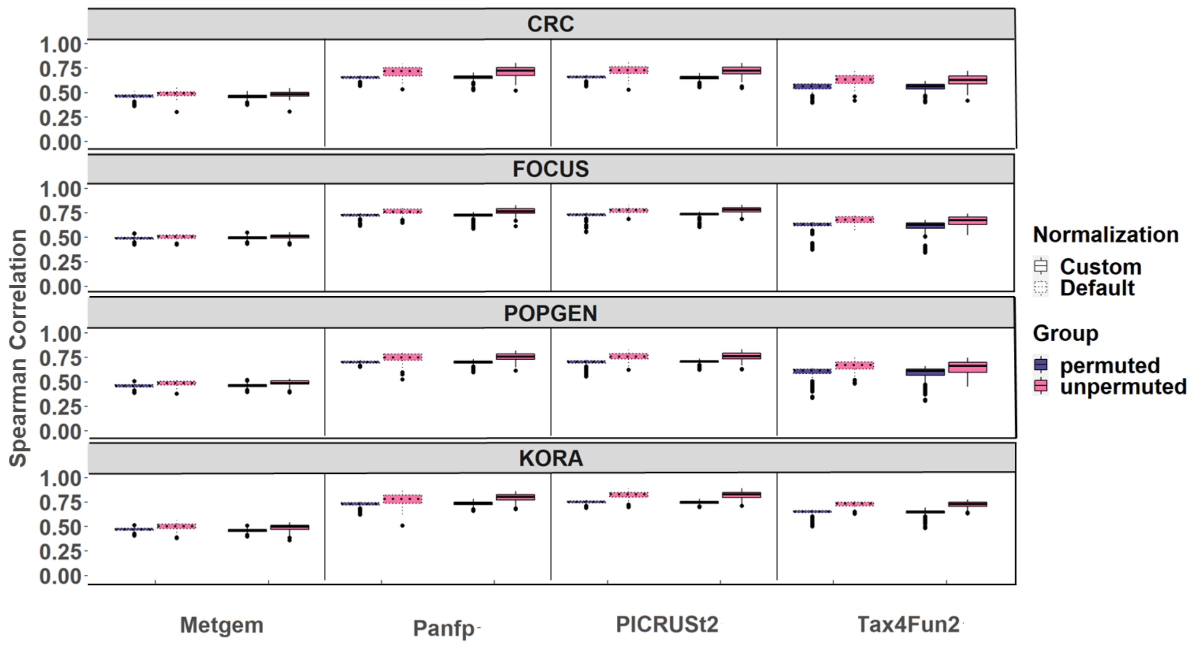


**Figure S1: Spearman correlations between metagenome predictions and shotgun metagenome sequencing in unpermuted and permuted datasets.** Validation of functional prediction tools comparing metagenome prediction performance against gold-standard shotgun MGS. Spearman correlations of gene composition estimated from metagenome sequencing and predicted with PICRUSt2, Tax4Fun, PanFP and MetGEM with default and customized normalization in unpermuted (blue) and permuted data (red) in all datasets. In each of the 100 permutations, every gene's abundance was permuted across samples independently.

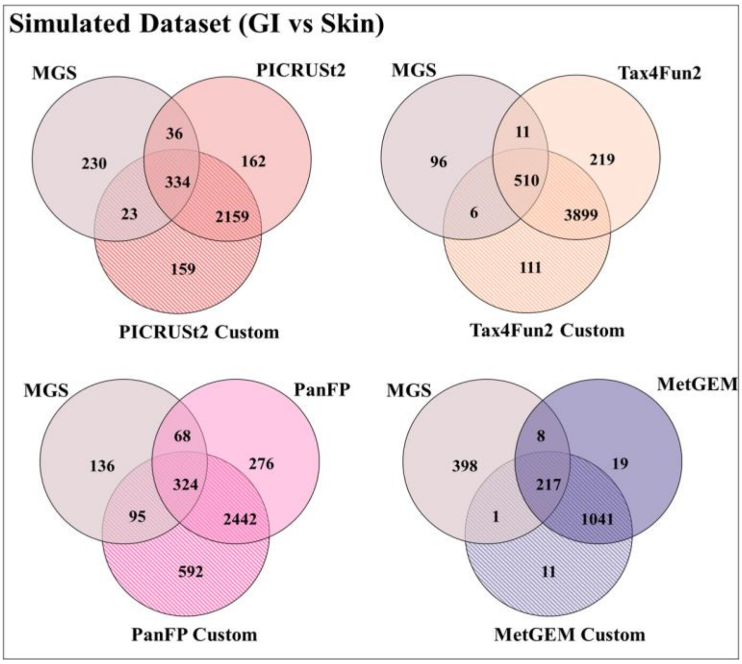


**Figure S2: The overlap of significant KO terms between different functional inferred tools and MGS results in the Popgen cohort. Overlapping significant KO terms can be used as an indicator of the accuracy of the predictions quantitatively.**

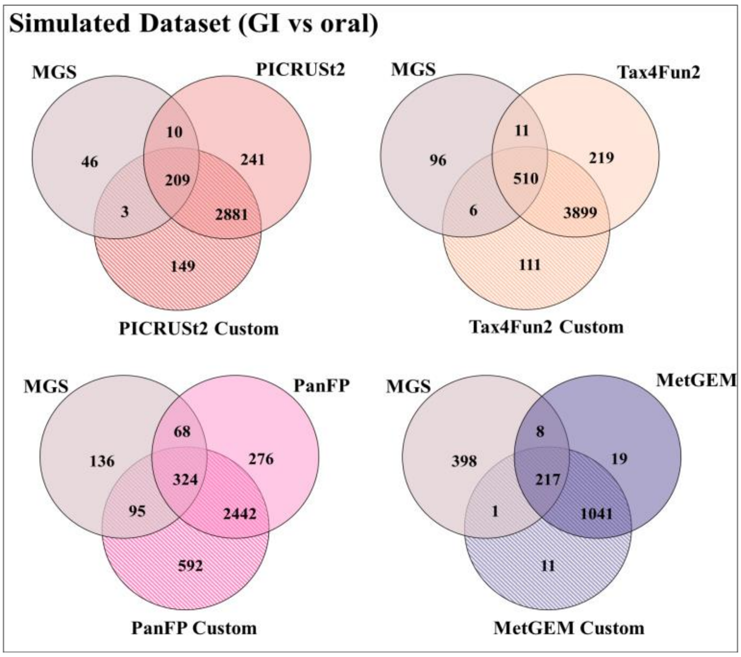


**Figure S3: The overlap of significant KO terms between different functional inferred tools and MGS results in the simulated datasets between GI and oral . Overlapping significant KO terms can be used as an indicator of the accuracy of the predictions quantitatively.**

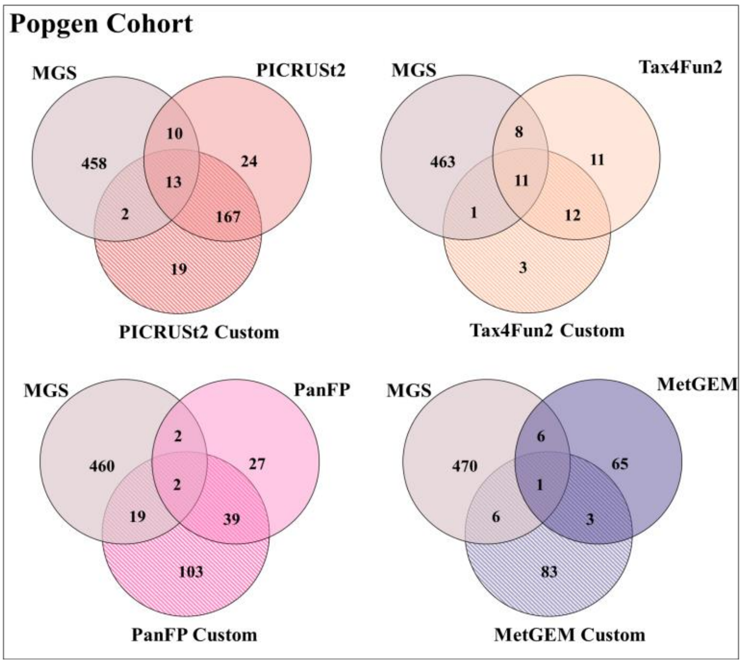


**Figure S4: The overlap of significant KO terms between different functional inferred tools and MGS results in the Popgen cohort. Overlapping significant KO terms can be used as an indicator of the accuracy of the predictions quantitatively.**

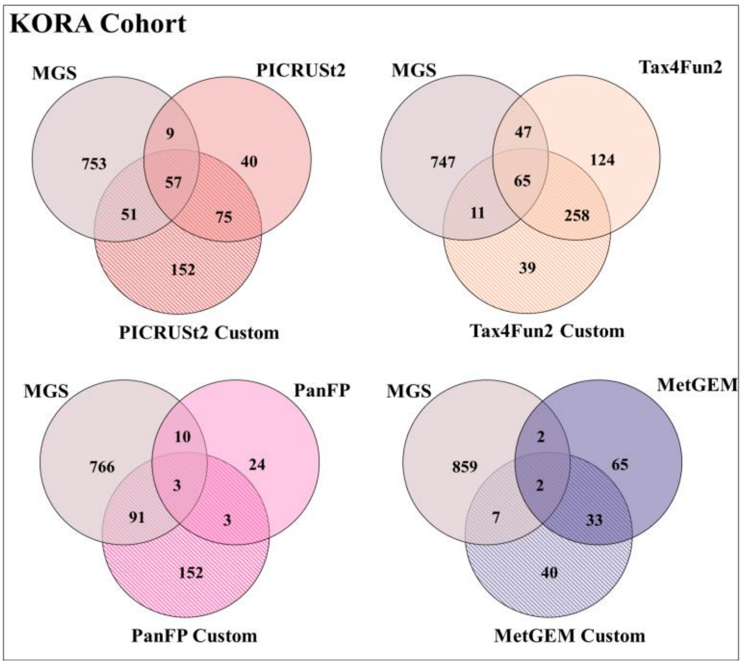


**Figure S5: The overlap of significant KO terms between different functional inferred tools and MGS results in the KORA cohort. Overlapping significant KO terms can be used as an indicator of the accuracy of the predictions quantitatively.**

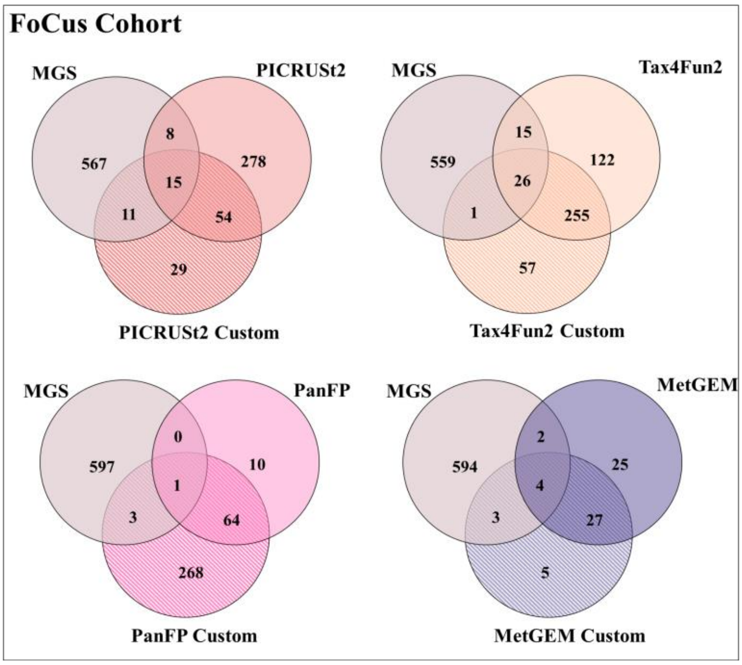


**Figure S6: The overlap of significant KO terms between different functional inferred tools and MGS results in the FOCUS cohort. Overlapping significant KO terms can be used as an indicator of the accuracy of the predictions quantitatively.**

### S2. The performance of functional inference tools varies widely across different KEGG functional categories

The results above suggest that 16S-based inference tools can recapitulate the general microbial environment and related microbial functions comparably well. We next investigated if prediction performance varies across functional categories, suspecting that some categories may be easier to predict than others. In general, KO terms are organized into a hierarchical structure, with each term belonging to a KEGG higher-level category using categorize_by_function.py provided by PICRUS2. For example, some categories may be more well-defined or have more established biological functions, making it easier to accurately predict the presence or abundance of related KO terms. Conversely, other categories may be more complex or poorly understood, leading to poorer prediction performance. To investigate whether prediction performance varies across functional categories, we performed an analysis that compares the accuracy of predicted KO terms within each KEGG higher-level category. By doing so, we can identify which functional categories are particularly challenging to predict, and whether there are any trends or patterns in the data that could explain these difficulties. This information could be useful for refining predictive models and improving the accuracy of predictions overall.

Level 1 KEGG functional categories group together related pathways and other functional annotations based on higher-level functions. Level 2 KEGG functional categories further subdivide the level 1 categories into more specific functional categories. For example, the level 2 category "Carbohydrate metabolism" is a subset of the level 1 category "Metabolism". The

KO terms themselves are organized into level 3 functional categories, which provide even more specific information about the functions of individual genes or gene products. Hence to evaluate the performance of functional inference tools at different levels of resolution, such as KEGG level 1, 2, and 3 functional categories, the predicted KO term abundances were aggregated at each level of the hierarchy using picrust1_categorize_by_func.R script. The categorization was based on a legacy table of mappings from KOs to BRITE hierarchy, which was used as a reference. The BRITE (Biomolecular Reaction/Interaction Database) [50] hierarchy is a hierarchical classification system used in the KEGG (Kyoto Encyclopedia of Genes and Genomes) database. It categorizes biological entities, such as genes, proteins, and compounds, based on their functional and interaction properties.  and compared to the corresponding abundances obtained from metagenomic sequencing data.  The categorization was based on a legacy table of mappings from KOs to BRITE hierarchy, which was used as a reference. Once the KO terms were aggregated at the level 1 KEGG functional categories, the Wilcoxon rank-sum test was performed to test for significant differences in KO abundances between the disease and healthy groups in each of the cohorts.

To analyze the consistency of functional inference tools and metagenome sequencing, the KO terms were combined into level 2 and level 3 KEGG functional categories as mentioned in the method section. We then sought to identify disease-associated functions through a Wilcoxon rank sum test for all level 2 and level 3 categories. We evaluated the results of inference methods considering significant changes observed in MGS data as ground truth using the F1, recall and precision score. Interestingly, we found that, except for the CRC cohort (Figure 5), functional profiling tools did not identify any true positives, indicating that they did not identify any significant functional pathways related to the disease state in the other cohorts (KORA, POPGEN, and FOCUS). Cancer-associated dysbiosis might lead to more substantial changes in the metagenome compared to other diseases which could be better reflected in the 16S rRNA data. However, further data are needed to corroborate this.

PICRUSt2 and Tax4Fun2 showed good performance at the level 3 functional categories. They rely on reference-based approaches to infer functional pathways in microbial communities which may be beneficial for detecting pathways at level 3. Metgem struggled at level 2 and level 3 functional categories. PanFP, which uses a pan genome-based approach, may be more effective at predicting functional pathways at level 2 because it does not rely on reference genomes or genome annotation data but instead uses a database of pangenomes from multiple species to predict metabolic pathways. This approach may be more flexible and better able to handle the diversity of microbial communities in different environments. Throughout our results we consistently noted that there was no discernible positive effect when employing customized copy number normalization techniques.

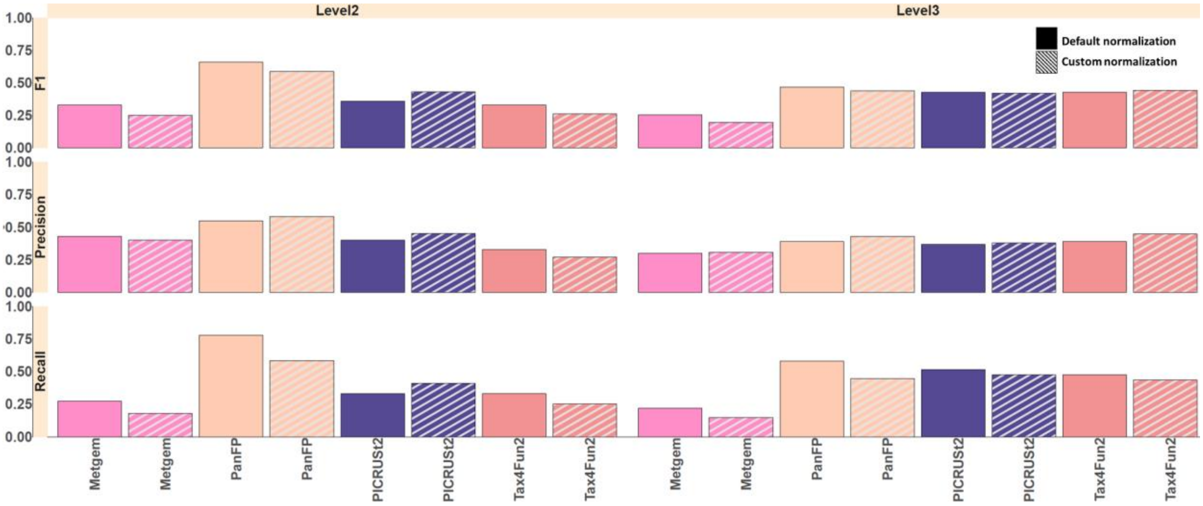


**Figure S7: Comparison of F1 score, recall, and precision for predicting the occurrence of KEGG functional categories in CRC cohorts versus MGS data, across three KEGG functional category levels.**

We performed differential abundance testing of KO terms using the Wilcoxon rank-sum test at different KEGG levels. As a first step, correlation of p-values from predicted and MGS-derived level 1 KEGG functional categories was performed. Different biological processes and pathways can have distinct features and complexities that may affect the performance of functional inferred tools. For example, metabolic pathways involve a complex network of reactions and regulatory mechanisms that may be different from those involved in signal transduction pathways. As a result, functional inferred tools that perform well in predicting functions in one category may not perform as well in another category. By evaluating the performance of functional inferred tools across different KEGG hierarchical categories, researchers can identify the strengths and limitations of each tool in predicting functions in different biological contexts. This can help guide the selection of appropriate tools for specific research questions and aid in the interpretation of functional genomics data. Moreover, the KEGG hierarchical categories provide a standardized classification system for gene functions, allowing researchers to compare the performance of different functional inferred tools in a consistent manner. This can facilitate the development and improvement of functional inferred tools and contribute to the advancement of functional genomics research. In this setting, metagenome inferred tools missed a large set of genes that are predicted by metagenome and likewise predicted many genes that are actually not found to be differentially abundant in MGS data.

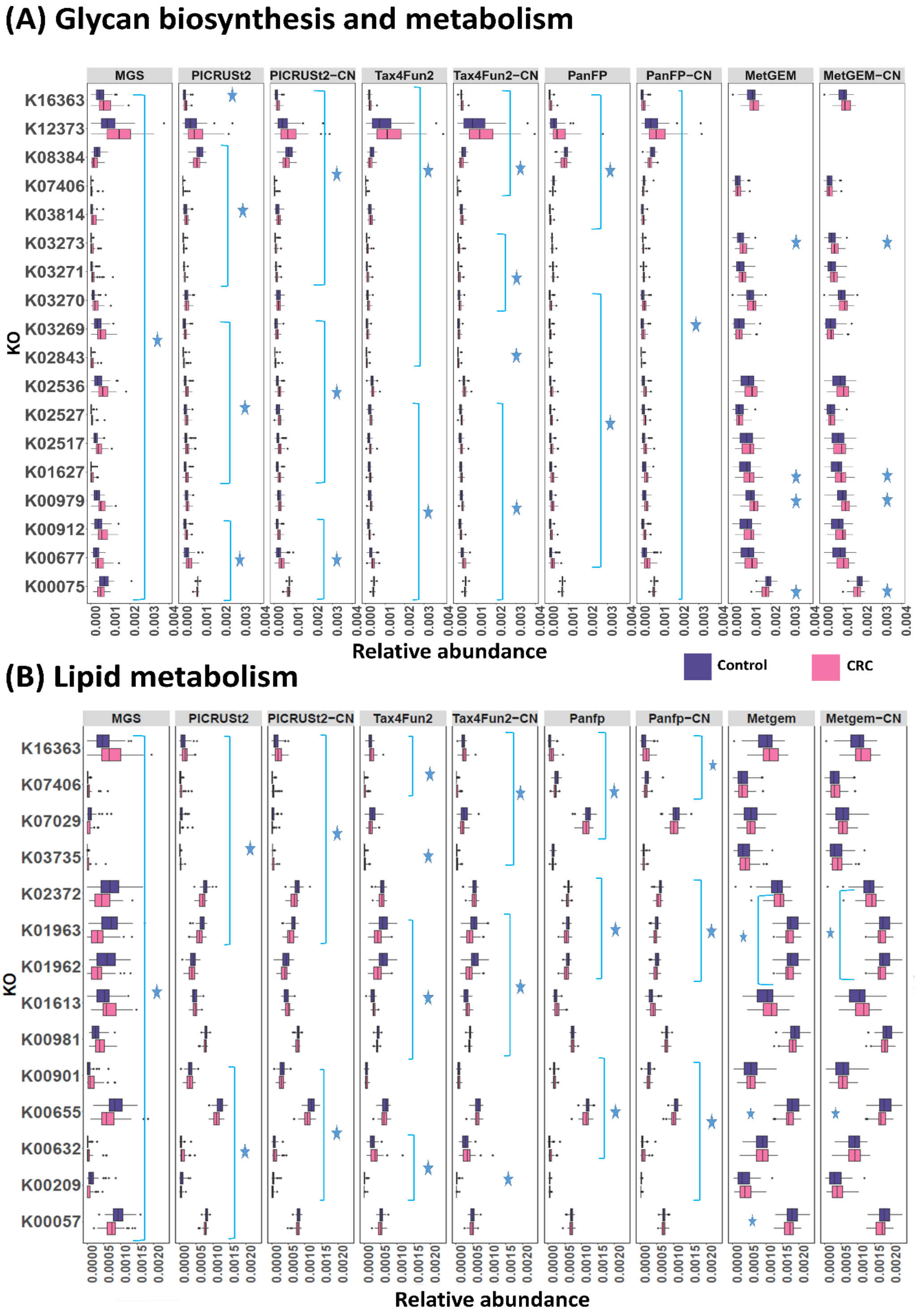


**Figure S8: Comparison of relative abundance distributions among major KEGG gene categories such as (A) glycan biosynthesis metabolism and (B) lipid metabolism between functional profiles produced by inference tools and those derived from MGS in the CRC cohort. A Wilcoxon rank-sum test was performed to compare the relative abundance between healthy and diabetes groups. The threshold of a p-value of less than 0.05 indicated a significant difference (*).**

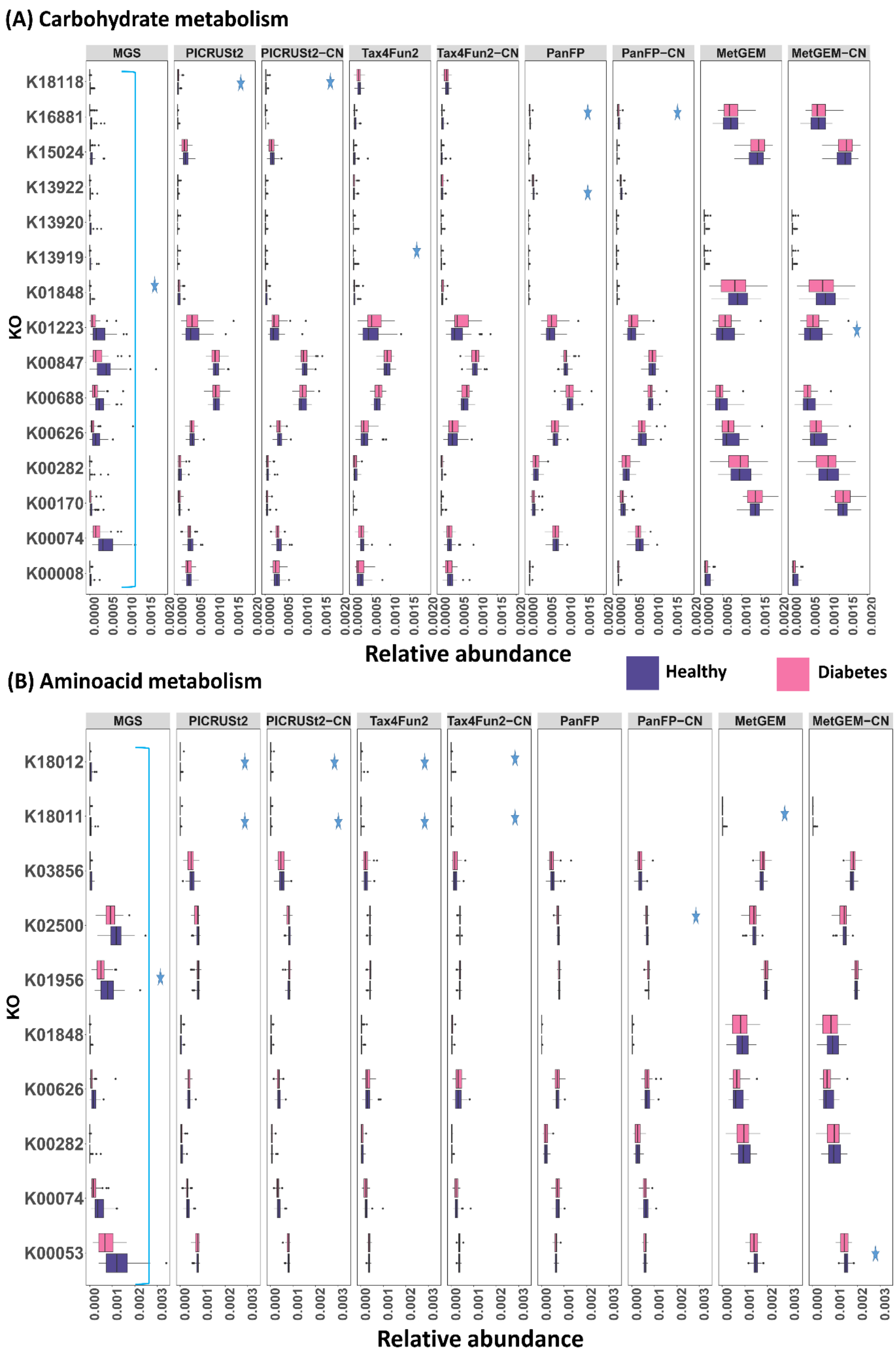


**Figure S9: Comparison of relative abundance distributions among major KEGG gene categories such as (A) carbohydrate metabolism and (B) amino acid metabolism between functional profiles produced by inference tools and those derived from MGS in the KORA cohort. A Wilcoxon rank-sum test was performed to compare the relative abundance between healthy and diabetes groups. A p-value of less than 0.05 indicated a significant difference (*).**

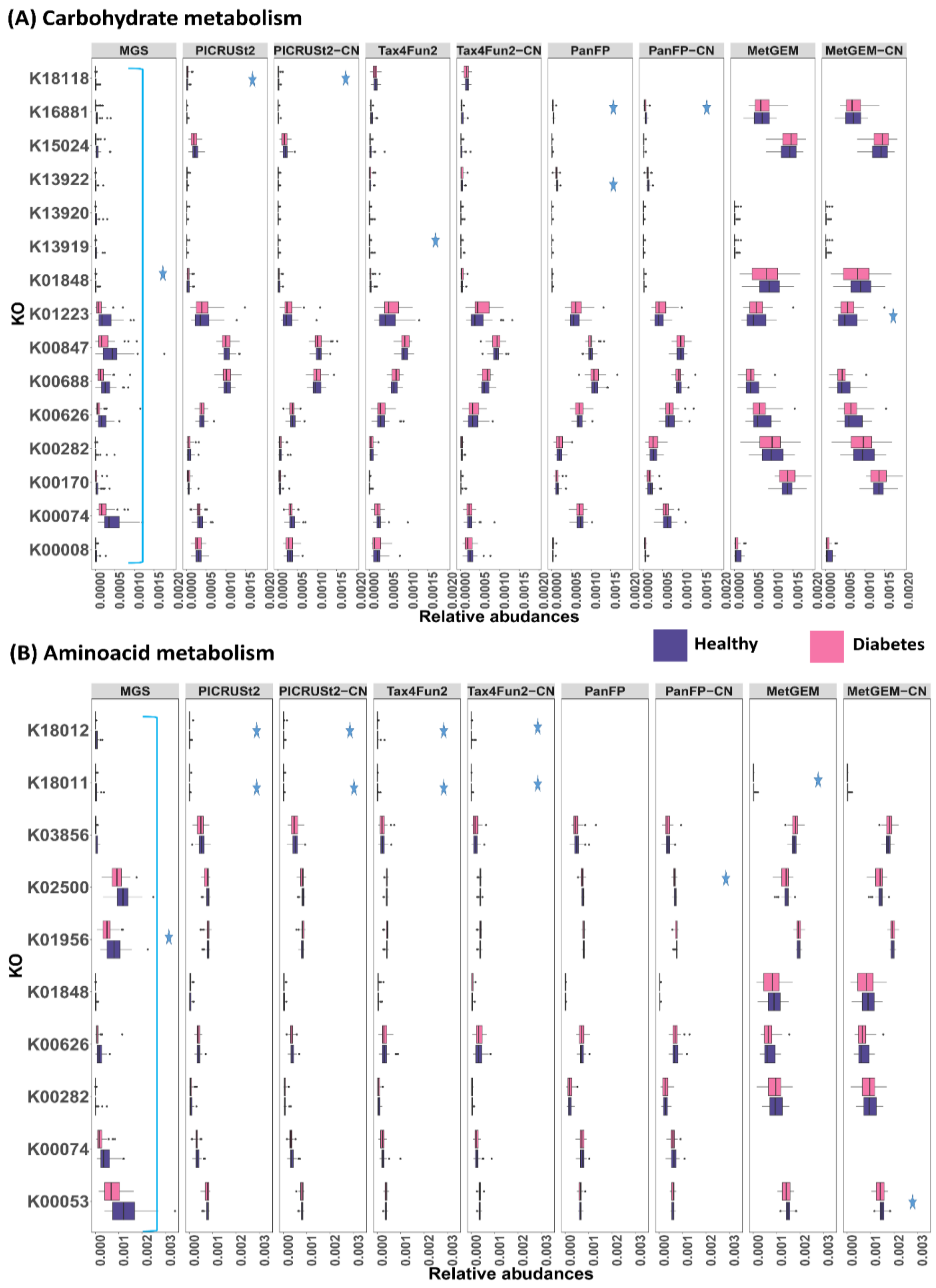


**Figure S10: Comparison of relative abundance distributions among major KEGG gene categories such as (a) carbohydrate metabolism and (b) amino acid metabolism between functional profiles produced by inference tools and those derived from MGS in the PopGen cohort. A Wilcoxon rank-sum test was performed to compare the relative abundance between healthy and obese groups. A p-value of less than 0.05 indicated a significant difference (*).**

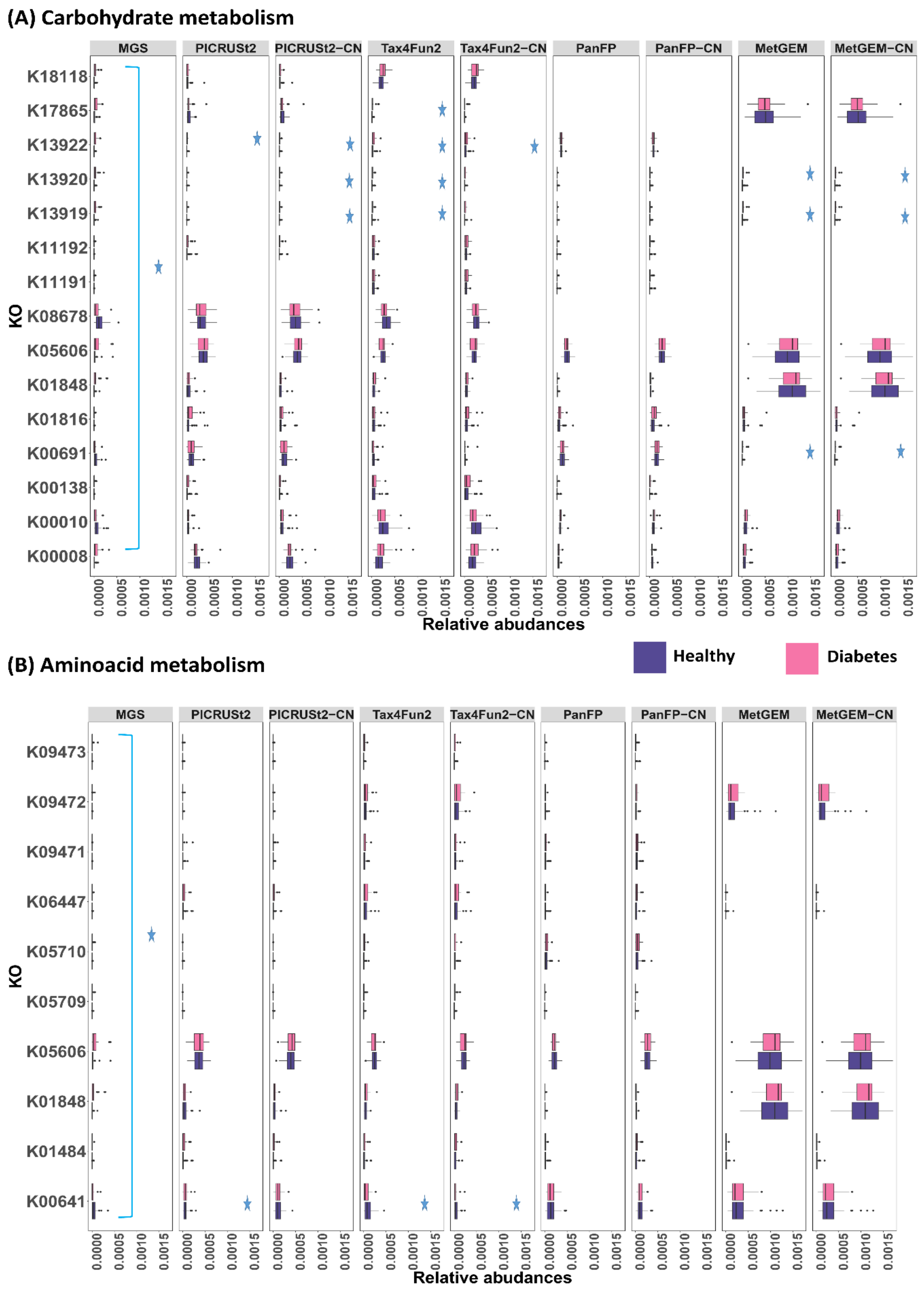


**Figure S11: Comparison of relative abundance distributions among major KEGG gene categories such as (a) carbohydrate metabolism and (b) amino acid metabolism between functional profiles produced by inference tools and those derived from MGS in the FoCus cohort. A Wilcoxon rank-sum test was performed to compare the relative abundance between healthy and obese groups. A p-value of less than 0.05 indicated a significant difference (*)**
